# Quantitative mapping of triacylglycerol chain length and saturation using broadband CARS microscopy

**DOI:** 10.1101/463000

**Authors:** A. Paul, Y.J. Wang, C. Brännmark, S. Kumar, M. Bonn, S. H. Parekh

## Abstract

Lipid droplets (LDs) are highly dynamic organelles that store neutral lipids, primarily triacylglycerols (TAGs), and are found in many cell types. While their primary function is to store excess energy, LDs are also modified in different disease states and during developmental processes. In many cases, not only the presence, but also the composition, of LDs can be equally important. In humans, LD composition has been linked to diseases such as type 2 diabetes; in plants and yeast, LD composition is relevant for engineering these organisms into biological factories in, *e.g.*, algal bioenergy or food oil production. Therefore, lipid analysis of biological LDs with high speed and accuracy *in situ* is a very active area of research. Here we present an approach for *in situ*, quantitative TAG analysis using label-free, coherent Raman microscopy to decipher LD TAG composition in different biochemically complex samples. Our method allows direct visualization of inter-LD compositional heterogeneity of physiological quantities – TAG chain length and number of C=C bonds – with sub-micrometer spatial resolution within 5-100 milliseconds. Combined with virtually no sample preparation, this approach should enable rapid and accurate TAG LD analysis for a variety of applications.

## Introduction

Many cell types use lipid droplets (LDs) as efficient storage depots to provide energy in times of scarce nutritional supplies. In order to build up these storages, free fatty acids, which are toxic to most cells, are esterified into neutral lipids, such as triacylglycerol (TAG) by acyl transferase enzymes. These hydrophobic TAG molecules form droplets together with other esterified molecules inside the cytosol and the droplets are covered by an emulsifying monolayer of phospholipids (1, 2). Additionally, various proteins are associated to LDs, *e.g*., the perilipin proteins in mammals or oleosins in plants, which can have various functions from anchoring lipases to serving as scaffolds for signaling molecules (2). LDs have been found across many different species with the general consensus being that LDs contain mostly TAGs and sterol esters, and to a lesser extent diacylglycerol. The ratios between these molecular species can vary vastly between different cell types and potentially for the same cell in different cellular states, *e.g.*, during diseases (3, 4). In mammals there is a specific cell type, adipocytes, designated to acquit lipotoxicity and store TAGs for energy usage. Outside of adipocytes, LDs also appear in many other mammalian tissues, especially when the mammal is exposed to high levels of fatty acid intake; such ectopic lipid accumulation has been identified to cause dysfunction and death of the lipid-laden cells. One example is in pancreatic ß-cells where excess lipid accumulation has been found to interfere with insulin secretion (5) and also lead to apoptosis (6).

Similar to mammals, prokaryotic yeast have lipid accumulations – largely full of TAGs – that allow cells to deal with lipotoxicity (7). TAGs in yeast can also be a desired product in the search for replacement of plant-based resources such as cocoa butter (8). Not only cocoa seeds, but many other plant seeds store TAGs in oil bodies (3), which are analogous to the lipid droplets in mammalian cells in that they are coated with amphipathic lipids and regulatory proteins (9).

Over the last 15 years it has become clear that LDs in a single organism can be divided into different subpopulations according to lipid composition and proteins in the LD phospholipid coating (10–12).

Moreover, the size and location of the droplets differ between cell types and organisms. A clear example is again mammalian white adipocytes where a large, central lipid droplet fills almost all of the cellular volume (13), whereas in, *e.g.*, muscle myocytes LDs appear smaller and dispersed in the cells (14–16). This same appearance of small LDs is also observed for drosophila fat bodies (17, 18) as well as in yeast (19, 20) and plant cells (21). In addition to different size, LD composition can also differ among individual cells (22) and even within single cells (23). Importantly, fatty acid composition, particularly carbon saturation level found in ectopic lipid droplets in skeletal muscle and chain length of free fatty acids in the blood, are known to correlate with increased insulin resistance (24) and stimulation of insulin secretion (25), respectively. Moreover, exposing cultured cells to saturated fatty acids is known to be cytotoxic whereas an equivalent concentration of unsaturated fatty acids can be relatively benign (26). Since fatty acids in cells are eventually stored in LDs, there is a need to quantify the lipid chemistry at the individual LD level in cells.

Traditional approaches for the characterization of lipids in cells or tissues involve the extraction of all lipids followed by derivatization and analysis with gas chromatography mass spectrometry (GC-MS) (27). The location of cellular LDs can be determined within cells using microscopy *via* fluorescence markers (28, 29) or label-free vibrational imaging such as coherent anti-Stokes Raman scattering (CARS) microscopy (30). In order to achieve spatial information coupled to chemical structures, three main methods have emerged during the recent years: (i) matrix-assisted laser desorption/ionization mass spectrometry imaging (MALDI-MSI), (ii) magnetic resonance imaging (MRI), and (iii) hyperspectral Raman or CARS microscopy. MALDI-MSI has been used to identify lipid species with high accuracy and > 1.4 μm spatial resolution in eukaryotic systems, *i.e.*, mouse brain and *paramecium caudatum* (31). This technique relies however on thin biological sections, which are embedded into an optimized matrix. Due to this extensive sample preparation and the relatively long measurement times, MALDI-MSI is slow in throughput. Additionally, freeze-thaw cycles as required for sectioning and subsequent embedding can cause LD mobilization and flowing in the tissue. MRI is well suited to visualize oil bodies *in vivo* from plants to humans (32, 33). With a resolution of around 10 μm in xy and 100 μm in z direction and the capability to image centimeter scale samples, this technique is applicable to investigate the structure of TAGs in larger lipid depots in entire organisms (34), but it is not readily equipped to spatio-chemically resolve fine structures, i.e. individual LDs. Vibrational spectroscopy, *e.g.*, spontaneous Raman and CARS, uses intrinsic molecular vibrations to visualize different macromolecular species: all lipids, all nuclear acids, or all proteins and can be used both *in vitro* and *in vivo* (30, 35). CARS microscopy relies on a resonant enhancement via nonlinear excitation to achieve higher signal strengths compared to spontaneous Raman scattering (for highly concentrated species), and the blue-shifted signal light is readily separated from linear fluorescence background. Using a hyperspectral approach, chemical information can be mapped with sub-μm resolution with complete vibrational spectra at each spatial location (36). From these spectra, ratiometric quantities have been employed to map TAG saturation in mammalian cells (37, 38) and in algae (39). However, these analyses are not equipped to study TAG chain length in combination with saturation. For that purpose, a least squares decomposition of single point Raman spectra from intracellular droplets has been employed by Schie *et al.* (40) to track the uptake of oleic and palmitic acid. Similarly, a factorization approach of hyperspectral CARS data has been used to study lipid dynamics in human stem cells (41). The latter two approaches focus on quantifying the amount of a small subset of known fatty acids that were supplied to the culture medium and ultimately incorporated in the LDs; however, it is not possible to use these methods determine the chain length and saturation of TAGs in native (unstimulated), single LDs.

In this work, we employ broadband hyperspectral CARS (BCARS) microscopy combined with a least squares decomposition to map the average number of C=C double bonds and chain length in TAGs in native LDs with sub-micrometer spatial resolution and ~ 5 − 100 millisecond acquisition times. We demonstrate applicability of this approach in artificial emulsions, microalgae, yeast, and mammalian cells, where our results compare favorably with mass spectrometry results. This analytical methodology should be readily transferable to other experimental setups and provides meaningful numbers that can be understood in a physiological context.

## Materials and Methods

### Materials

TAG and oil standards, and BSA were purchased from Sigma-Aldrich (Darmstadt, Germany): glyceryl trioctanoate (8:0, T9126, CAS 538-23-8), glyceryl tripalmitate (16:0, T5888, CAS 555-44-2), glyceryl tripalmitoleate (16:1, T2630, CAS 20246-55-3), glyceryl tristearate (18:0, T5016, CAS 555-43-1), glyceryl trioleate (18:1, T7140, CAS 122-32-7), glyceryl trilinoleate (18:2, T9517, CAS 537-40-6), glyceryl trilinolenate (18:3, T6513, CAS 14465-68-0), canola oil (46961, CAS 12096-03-0), cottonseed oil (47113, CAS 8001-29-4), olive oil (47118, CAS 8001-25-0), peanut oil (47119, CAS 8002-03-7), soybean oil (47122, CAS 8001-22-7), sunflower seed oil (47123, CAS 8001-21-6), and BSA (05470, CAS 9048-46-8). Long chained TAGs were purchased from Larodan (Solna, Sweden): glyceryl triarchidonoyl (22:4, 33-2040, CAS 23314-57-0), glyceryl tridocosahexaenoyl (22:6, 33-2260, CAS 124596-98-1).

### Pure and diluted oils

Pure oils were measured either at room temperature (8:0, 16:1, 18:1, 18:2, 18:3, 20:4, 22:6) or at 86°C (16:0, 18:0) with a homebuilt Peltier temperature stage. Oils diluted in CCl_4_ and toluene-d8 were measured at 10°C.

### Stabilized emulsions

Oils were encapsulated at 10 vol.% in 3 wt.% agarose with low gelling temperature after thorough vortexing and sandwiched between two glass cover slips (#1). Supplemental Fig. S1 *a* and *b* shows that different sized lipid droplets are formed this way. Due to the vortexing, water -oil-water double emulsions can form (droplets with dark spots in Supplemental Fig. S1 *b*). They were excluded from this study.

### Cell culture

3T3-L1 adipocytes were cultured on glass-bottom petri-dishes and differentiated as previously described (42, 43). RAW 246.7 and HEK-293 cells were cultured on glass and grown in medium supplemented with 2mg/ml low density lipoprotein for 24 and 48 hours, respectively. Before imaging, all cells were fixed in 4 vol.% paraformaldehyde, crosslinking proteins without effect on lipid chemistry. CENPK 113-11C *Saccharomyces cerevisiae* and *Yarrowia lipolytica*, TAG producing yeast strains, were cultured as previously described (8, 44) and fixed in paraformaldehyde. *Nannochloropsis* sp. (CCAP211/78) was cultivated according the methods used for strain maintenance and flask cultivation detailed in Mayers *et al.*(45). Cultures were allowed to deplete media nitrogen for the purposes of this analysis to generate cells with higher lipid content. Both yeast and algae cells were subsequently encapsulated in 3 wt.% agarose with low gelling temperature, and sandwiched between two cover slips. An overview of all used samples can be found in Supplemental Fig. S1 *c-l* showing the morphology of the different cell types as well as their intracellular lipid droplets.

### Coherent anti-Stokes Raman scattering microscopy

Lipid droplets in artificial emulsions, 3T3-L1 adipocytes, yeast, and algae were visualized with CARS microscopy. A specific molecular vibration, *e.g.*, the symmetric CH_2_ stretching at 2845 cm^−1^, is targeted with the field generated by two laser pulses (with energies ω_pump_ and ω_stokes_) overlapped in time and space. A third pulse (ω_probe_) is then used to generate the CARS signal (ω_anti-Stokes_ = ω_pump_ + ω_stokes_ - ω_probe_). The CARS imaging was performed with a SP 5 II TCS CARS microscope (Leica, Mannheim, Germany) equipped with a pico-second pulsed laser source at 817 and 1064 nm (APE, Berlin, Germany) using a water immersion objective (L 25x, NA=0.95, Leica) to achieve tight focusing conditions. CARS signals were collected in forward direction with a Leica CARS 2000 filter set in front of the photomultiplier tube. For 3T3-L1 adipocytes autofluorescence was collected in the epi direction using the Leica FITC filter set.

### Broadband CARS microscopy

The home-built setup with custom software written in LabVIEW (National Instruments) has been described in detail previously (46). Briefly, from a dual-output laser source (sub-ns pulses, 32 kHz, Leukos-CARS; Leukos, Limoges, France) a super continuum Stokes beam is generated using a photonic crystal fiber (100 μW nm^−1^, 1050-1600 nm). This Stokes beam were then overlapped with the fundamental pump (and probe) beam at 1064 nm in the focus of an inverted microscope (Eclipse Ti, Nikon) equipped with a xyz piezo stage (Nano-PDQ 375 HS, Mad City Labs) for sample scanning. A near-infrared objective (100x, NA 0.85, PE IR Plan Apo, Olympus) was used to tightly focus the two beams onto the sample with the resulting CARS signals being collected in the forward direction by a 10X objective (M-10x, NA 0.25, Newport) for all samples except the mammalian cells, where a water dipping collection (W N-Achroplan 10x, NA 0.3, Zeiss) was employed for collection. The collected CARS signal was filtered through a notch (NP03-532/1064-25, Semrock) and short-pass (FES1000, Thorlabs) filter before being dispersed in a spectrometer (300 lines mm^−1^, 1000 nm blaze, Shamrock 303i, Andor) and detected on a deep-depletion CCD (Newton DU920P-BR-DD, Andor) with a spectral pitch of 4 cm^−1^/pixel. Images were collected with 0.1-0.5 μm step sizes in the xy plane and each pixel was illuminated for 5-100 ms.

### Hyperspectral data analysis

All data were analyzed with custom routines in IgorPro (v.6.37, WaveMetrics) and MATLAB (R2016a, MathWorks). First, the imaginary component of the third-order Raman susceptibility was retrieved with a modified Kramers-Kronig algorithm (47) with glass as reference followed by error phase correction using either (i) an iterative noise-maintaining approach (48), which is model-free with the use of a second-order Savitzky-Golay filter smooting filter over 101 spectral points (404 cm^−1^) for mixed samples or (ii) an iterative fourth-order polynomic function for pure samples (48). The resulting spectra are referred to as Raman-like (RL) in accordance with previous publications (36, 49). The spectral points between 1200 – 1786 cm^−1^ and 2828 – 3102 cm^−1^ were selected for all subsequent processing and fitting.

### Lipid chain length and number of C=C bonds maps

The fitting method has been adapted from Waschatko *et al*.(50) and applied on full hyperspectral images. All RL spectra were aligned and normalized, *i.e.*, arbitrary intensity of 1, to the 1440 cm^−1^ CH_2_ deformation peak. A training matrix was formed from the RL spectra of 8:0 TAG, 16:1 TAG, 18:1 TAG, 18:2 TAG, TAG 18:3, TAG 20:4, and a mixture of 50% 8:0 and 50% 18:1 (v/v) (spectra Supplemental Fig. 2). and 100 mg/ml BSA in MilliQ-water. A least squares decomposition with known covariance (Table 1, Fig. 1A) was used in order to generate three components representing the presence of TAGs (C_TAG_), the number of C=C bonds per chain (C_#C=C_), the number of CH_2_ per chain (C_#CH2_). The spectrum of 100 mg/ml BSA was used as a generic protein component (C_protein_). Further, a fifth component (C_dilution_) attributed to spectral changes upon dilution of TAGs was generated using a global fitting on spectra of diluted TAG 16:1 and 18:1 in CCl_4_ (Supplemental Fig. S3). Therefore four components were assumed: pure TAG 16:1, pure TAG 18:1, pure CCl_4_, and an unknown dilution component. This dilution component was then refined to best fit the seven diluted spectra.

**Figure 1.**
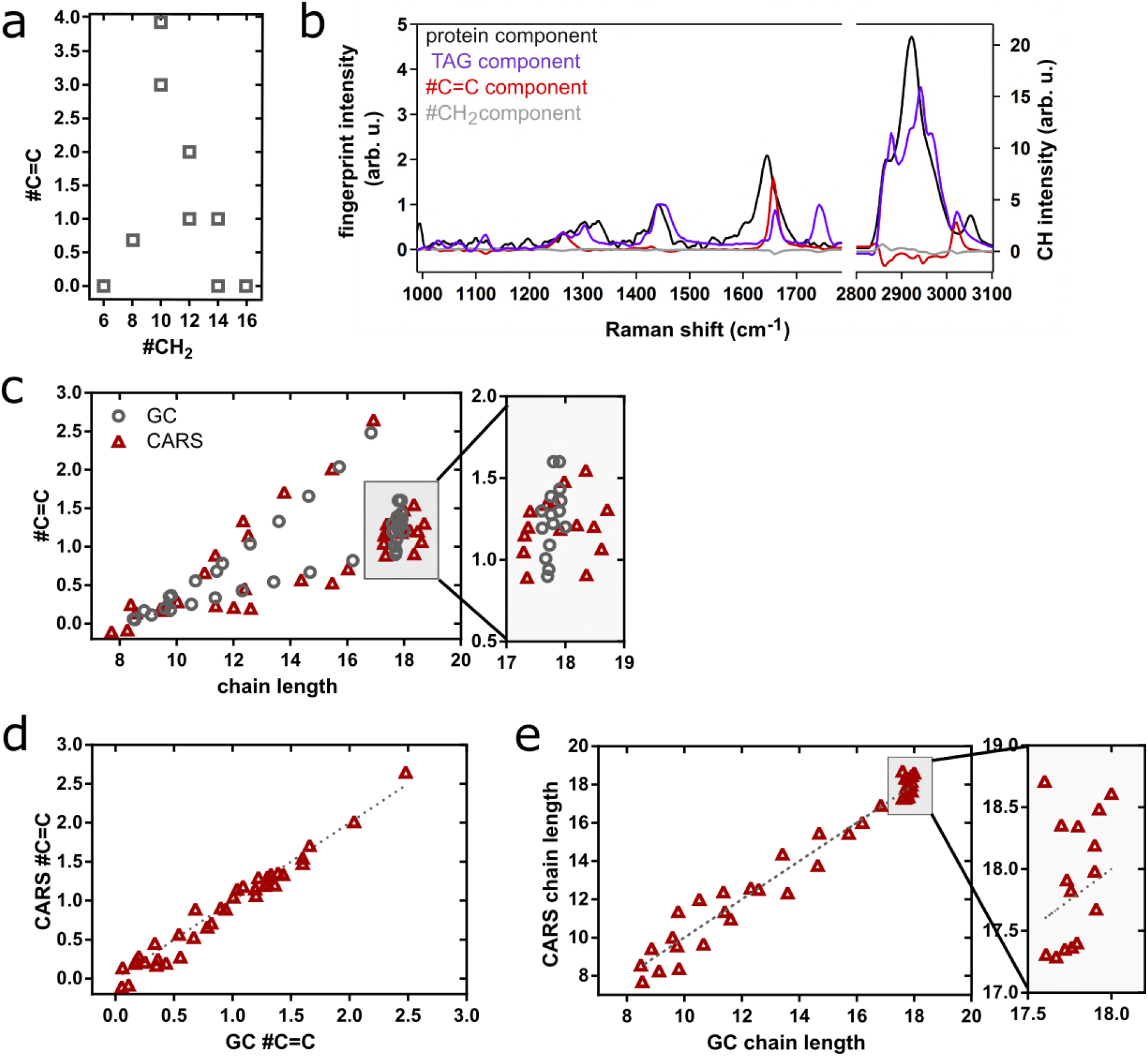
#C=C bonds and #CH_2_ groups for TAGs used in the training matrix to retrieve TAG, #C=C, and #CH_2_ components (*a*). The covariance table (Table 1) and spectra in *a* were used to produce three lipid components, which were used to recalculate experimental spectra. A generic protein component was also added for use in complex cellular LDs (*b*). A set of standard food oils, their mixtures, and TAG mixtures were analyzed and compared to the same quantities extracted from GC-MS measurements (c-e). The closer the red triangles (BCARS) are to the gray dots/line (GC-MS) the smaller the error. In d and e, dotted lines represent perfect agreement for the BCARS and GC-MS methods.

**Figure 2.**
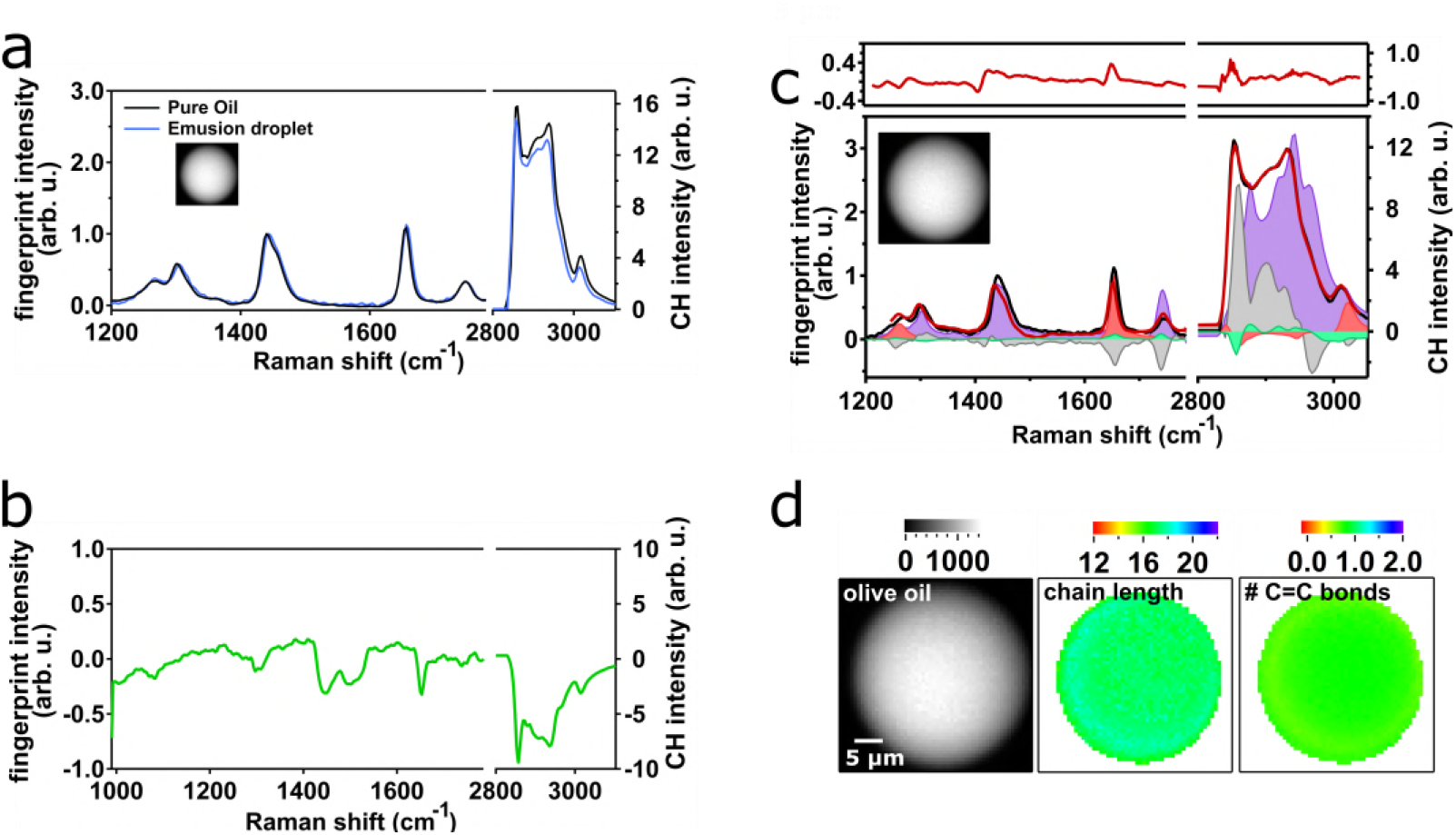
Normalized spectra (intensity of 1 at 1440 cm^−1^) of pure olive oil (black) and olive oil in the stabilized emulsion (blue) (*a*). Derived component representing the spectral differences caused by dilution of oils (*b*). Reconstruction (red line) of the experimental spectrum from the middle of the droplet (black line, normalized to 1440 cm^−1^) using the TAG (violet fill), #C=C (red fill), #CH_2_ (grey fill), and dilution (green fill) components (*c*). Fitting residuals are plotted as red line in the upper panel in *c*. Image of an olive oil emulsion droplet and computed maps for chain length and # C=C bonds (*d*) using the components from *c*.

**Table 1.** Known covariance for least squares decomposition.

| Molecule | TAG component | #C=C (per chain) | #CH<sub>2</sub> (per chain) |
| --- | --- | --- | --- |
|  |  | component | component |
| 8:0 TAG | 1 | 0 | 6 |
| 16:0 TAG | 1 | 0 | 14 |
| 16:1 TAG | 1 | 1 | 12 |
| 18:0 TAG | 1 | 0 | 16 |
| 18:1 TAG | 1 | 1 | 14 |
| 18:2 TAG | 1 | 2 | 12 |
| 18:3 TAG | 1 | 3 | 10 |
| 20:4 TAG | 1 | 4 | 10 |
| 50% 8:0 and 50% 18:1 (v/v) | 1 | 0.68 | 8.05 |

The component spectra as well as all spectra to be fitted were then interpolated with by a factor of 6 to ensure sub-pixel shifts were possible before fitting to a target spectrum with the following function:

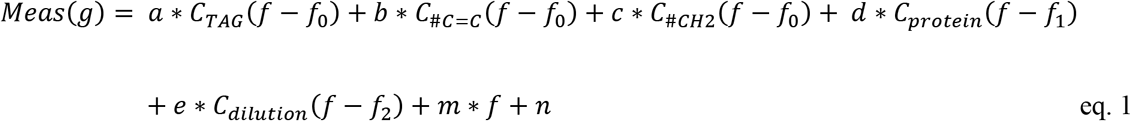

*a, b, c, d*, and *e* are weights of the five components, *f* is the Raman shift, and *f_0_*, *f_1_*, and *f_2_* allow for slight shifts of lipid, protein, and dilution components in the measured data (*Meas*). The values for *f_0_*, *f_1_*, and *f_2_* were always found to be less than 4 cm^−1^, less than the frequency spacing between adjacent CCD pixels (hence the necessity for interpolation), which is reasonable given the months timespan over which the data reported here was acquired. *m* and *n* allow for linear baseline offsets to be accounted for. For the spectral fitting of pure TAGs, oils and their mixtures only the three lipid components were used, for the emulsions the dilution component was added, and the cells the protein and dilution components were added. (all seed values: Supplemental Table S1). All weights for the components were constraint to be > − 0.1.

To recalculate the number of double bonds per chain and the average chain length of the TAG from the fit parameters, the weight of all components were divided by *a* to normalize *a* to 1, as was done in the training matrix. This allowed a read-out of the #C=C as *b/a*, #CH_2_ as *c/a*, and the chain length as 2*(#C=C)+(#CH_2_) +2 (accounting for end groups). For image analysis, a mask was generated based on the CH_2_ (2845 cm^−1^) intensity of the RL spectra in order to include only lipid spectra in the fitting, then a pixel-by-pixel fitting was done based on the method described above for single spectra. After the image fit, an additional threshold ensuring chain lengths were within 12-24 and #C=C > − 0.1 was applied to all images to eliminate unreasonable values

### Statistical testing

An unpaired two-tailed t-test with Welch’s correction was employed to analyze statistical significance between diameter, TAG chain length and saturation in the two adipocyte lipid droplet populations using GraphPad Prism 6.0. Values were ranked: P > 0.05 (ns, not significant), P ≤ 0.05 (*), P ≤ 0.01 (**), P ≤ 0.001 (***), and P ≤ 0.0001 (****).

## Results

### BCARS decomposition characterizes standard oils and TAG mixtures with ± 0.69 CH_2_ chain length and ± 0.15 #C=C accuracy

To establish a method for extracting average chain length and number of C=C from images of TAGs, we first produced a least squares decomposition with known covariances using a training matrix of TAGs. Our training matrix consisted of quantitative, linear Raman-like (RL) spectra from BCARS spectra (see Methods) of TAGs and their mixtures spanning the physiological range of fatty acid double bonds and chain lengths (Fig. 1 *a*, Table 1, full spectra in Supplemental Fig. S2). From the training matrix, three components could be extracted (Fig. 1 *b*): a TAG glycerol backbone component, the number of C=C per TAG, and the number of CH_2_ groups in the TAG chain. The TAG component represents the most common spectral features found in all TAGs in the training matrix while the number of C=C and CH_2_ components represent spectral changes that depend on the number of C=C double bonds or the chain length, respectively, as reflected in the known covariance (Table 1).

As our BCARS has a focal volume of 0.6 × 0.6 × 5 μm^3^, this method produces a molar average chain length and number of C=C bonds for all TAGs within this volume. The chain length discussed here refers the average number of carbons in one TAG chain, including the ester and CH_3_ end groups. Using the three lipid components, the number of C=C bonds, and the chain length from standard oils, mixtures of standard oils, and TAG mixtures was derived (Table S2 in the Supporting Material). The resulting parameters derived from the BCARS method were compared to the molar averages from GC-MS measurements to evaluate the error in our method. The overall agreement between the BCARS and GC-MS measurements, especially when both chain length and #C=C are taken into account, is clear (Fig. 1 *c*). On average our method is accurate for the number of C=C to within ±0.15 (Fig. 1 *d*) and for the chain length to within ±0.69 CH_2_ (Fig. 1 *e*). The error is evenly distributed around the GC-MS values, suggesting no systematic over-or underestimation.

### Imaging chain length and #C=C in oil emulsified droplets

Having established the ability to accurately quantify TAG chemistry in individual spectra, we next tested ability to map TAG chemistry in space. For that purpose we acquired images of olive oil droplets in oil-in-water emulsions to determine the accuracy of our method in a heterogeneous sample. Olive oil was mixed with melted agarose and vortexed. This sample preparation method generated oil-in-water emulsions but also more complex water-in-oil-in-water double emulsions (Supplemental Fig. S1 *b*). We chose this preparation method over a ‘cleaner’ approach to mimic the complexity of biological spectra, where several macromolecular species can be in the focal volume at once. Thus, we expect more complex spectra and indeed the emulsions showed a different spectral shape compared to pure oils (Fig. 2 *a*), similar to what we found in oil diluted in organic solvents. In order to account for this shape of the emulsified oil, we recorded spectra from a set of oils (18:1 and 16:1 TAGs) diluted in carbon tetrachloride (Supplemental Fig. S3) and used a global fitting approach to generate a spectral component that captured the spectral changes caused by dilution (Fig. 2 *b*). We validated this component against other diluted oils (Supplemental Table S3), which confirms the validity of the approach. The inclusion of this component allowed us to fit spectra from the olive oil emulsions and obtain C=C and chain length values within our error from pure oils (Fig. 2 *c,d*), even if there are small-scale impurities that effectively dilute the oil.

### Determining LD TAG chemistry *in situ* in biological organisms

The biological samples we studied had a broad range of LD sizes. *S. cerivisae* and *Nannochloropsis sp.* both have submicron sized lipid droplets (Supplemental Fig. S1). This LD size is small compared to their cellular volume, different from mammalian adipocytes, which can exhibit LDs that are several tens of microns in diameter and occupy the entire intracellular space. These submicron-sized lipid volumes can be challenging to analyze using standard methods like lipid extraction followed by GC-MS, where large culture volumes can be required, and cell sizes can approach the limit of mass spectrometry imaging. We also included *yarrowia lipolytica*, a yeast specialized for production of lipids as an alternative microorganism with larger LDs. For all these microorganisms, LDs have been identified label-free with single-frequency CARS microscopy of the symmetric CH_2_ vibration (at a frequency of 2845 cm^−1^) (51, 52), but it was not possible to use CARS to investigate the composition inside the droplets. As mammalian examples, differentiated 3T3-L1 adipocytes, LDL-fed RAW and LDL-fed HEK cells were investigated. As stated above differentiated 3T3-L1 cells have lipid droplets that can occupy nearly the entire cell (of order 10 μm) while LDL-fed RAW and HEK cells exhibit smaller, microsized LDs (Supplemental Fig. S1).

For these complex biological samples, a fifth component – the spectrum of BSA – was included to be able to account for a generic protein signal (Fig. 3 *a*), which will certainly be found in close proximity to LDs in these samples. This proximity will lead to spectra containing a mixture of protein and lipids at the LD edges at the very least, and possibly even those in the center of the LD for droplets that are smaller than the axial span of our excitation volume. Hyperspectral datasets from the different biological species were fit with all five components as described in the methods section, from which we produced images of carbon chain length and # or C=C bonds (Fig. 3 *d*). Immediately obvious from the images in Figure 3d are that the LDs in all biological systems are considerably more heterogeneous than in the stabilized emulsions, as expected. When averaging over the all images for each type of sample (aggregating from the single pixel basis), we found the average chain length in LDs spans from 16.7-18.8 over all species (Fig. 3 *b*), while more variety (compared to the average chain length over all species) is found in the # of double bonds, which spans from 0.29-0.94 (Fig. 3 *c*).

**Figure 3.**
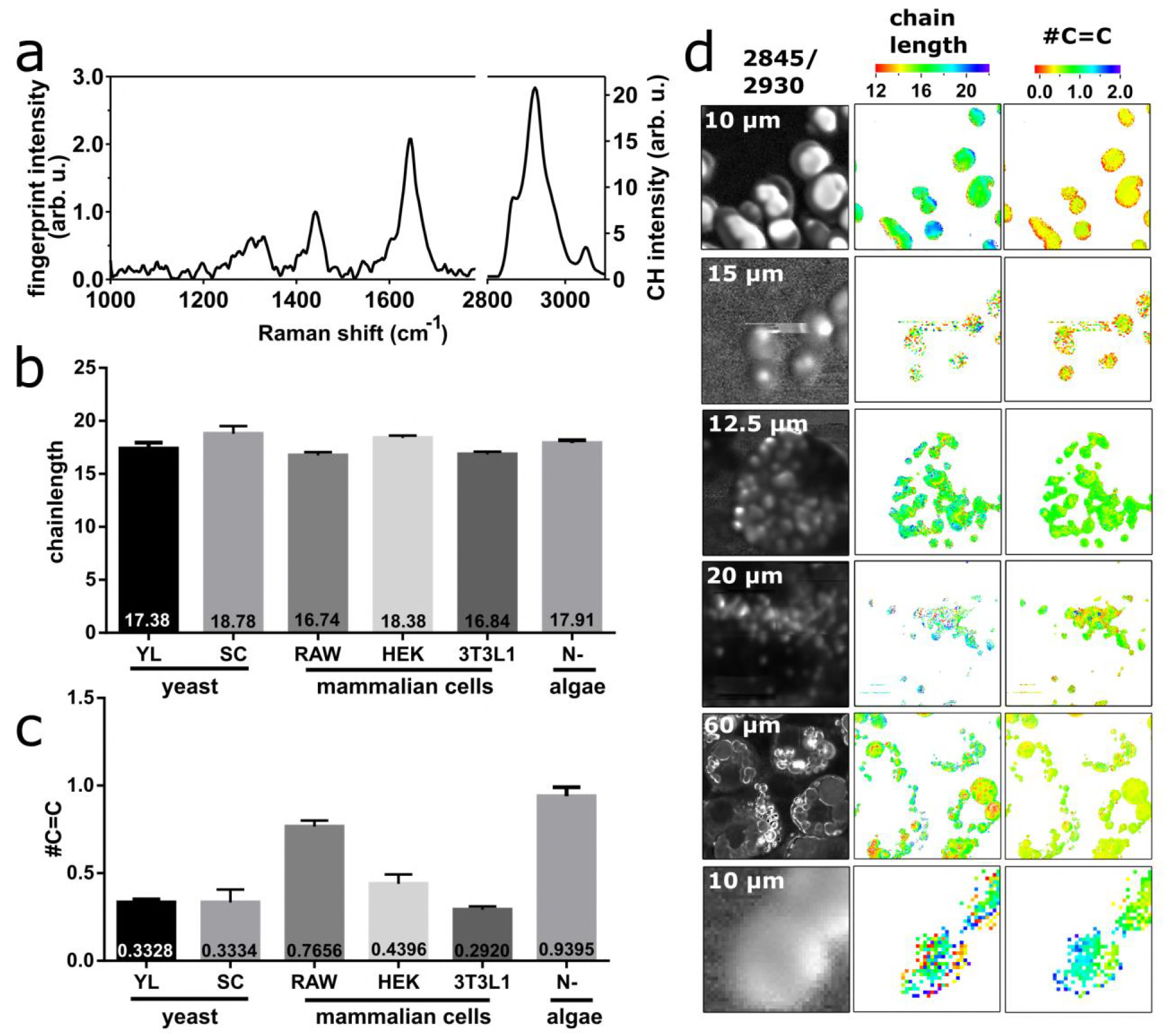
A generic protein component (BSA) was added as fifth component to improve the fitting of LD spectra from complex biological samples, where several macromolecule classes can be inside the focal volume (*a*). BCARS images from different biological species were analyzed and the average chain length (*b*) and number of double bonds (*c*) were determined by averaging all lipid containing pixels from multiple different cells in each category. Representative images displaying the cellular morphology (via the ratio of 2845 cm^−1^ over 2930 cm^−1^), chain length and #C=C distribution are shown in *d*. The species are displayed in the same order as in *b, c*. The number of analyzed images is: *yarrowia lipolytica* (YL) n=5, *S. cerivisae* (SC) n=5, RAW n=4, HEK n=3, 3T3-L1 n=27, and nitrogen starved *Nannochloropsis sp.* (N-) n=5. Values are displayed as mean ± SD.

The results from BCARS imaging of lipid chemistry for *yarrowia lipolytica*, representative of microoganisms, and 3T3-L1, representative of mammalian cells, were quantitatively compared to the GC-MS data from extracted neutral lipids (Supplemental Table S4 and Method S1). For bonds *yarrowia lipolytica* and 3T3-L1, we found average chain length / # of C=C bonds of 17.6/0.28 and 16.1/0.25, respectively, which is within our experimental error from the spatial averages shown in Figure 3c and 3d.

While all samples showed spatial variation, we were particularly drawn to that seen in the differentiated 3T3-L1 cells as there seemed to be two major classes of LDs in cells: 1) larger LDs that were fewer in number and 2) Smaller LDs that were quite numerous. Further, from looking at the images in Figure 3d, these LD classes seemed to contain different TAGs. Indeed, we found a weak inverse correlation between droplet diameter and chain length (Fig. 4 *a*) and between droplet diameter and number of double bonds (Fig. 4 *b*). If the droplets are split into two approximately equal groups according to their diameter, these differences become significant (Fig. 4 *c*). Small LDs (< 2 μm) have an average chain length of 17.8 with 0.35 double bonds, while big LDs have 16.7 and 0.31.

**Figure 4.**
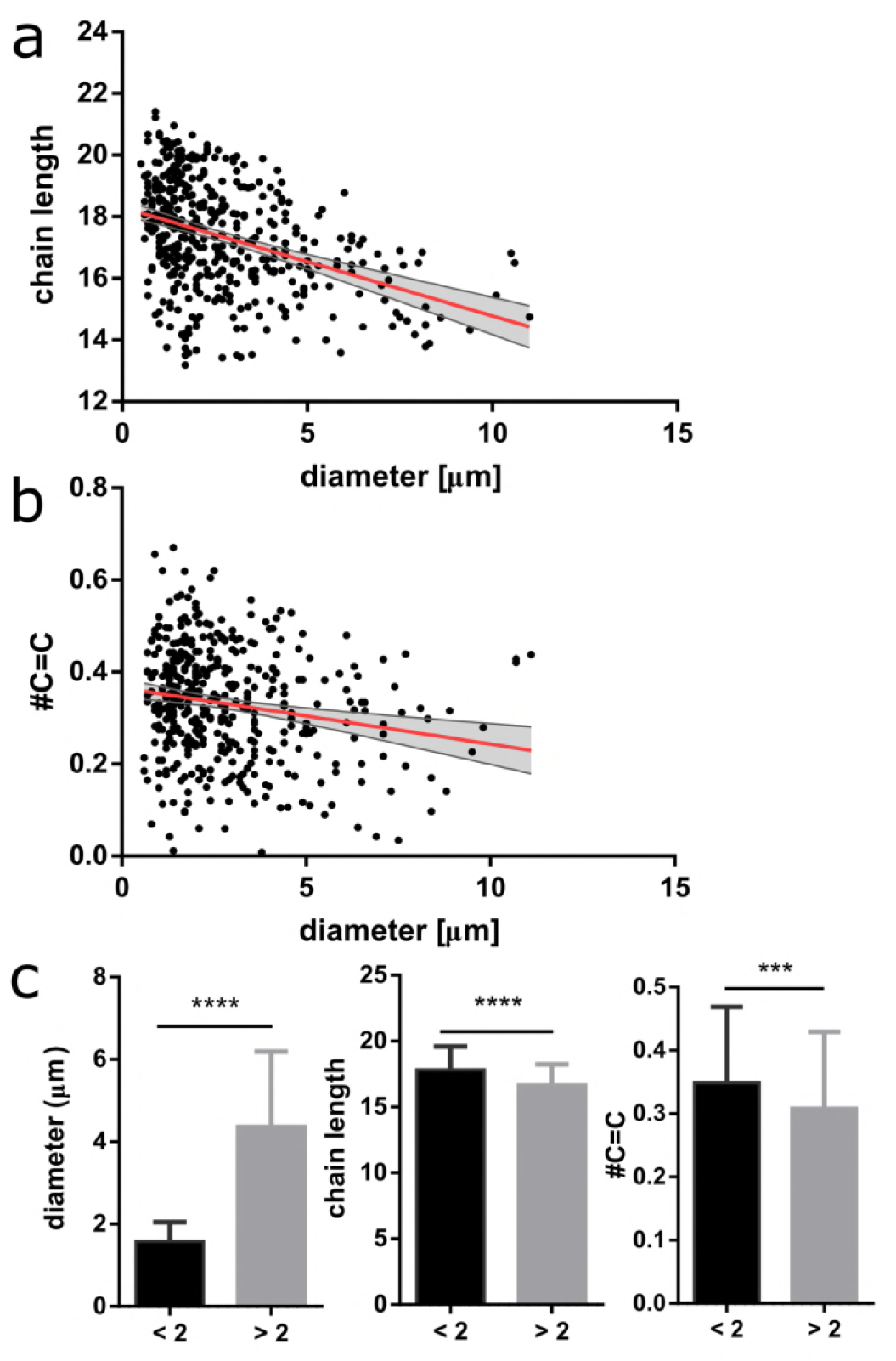
Chain length (*a*) and number of double bonds (*b*) show a negative correlation with droplet diameter. Splitting the droplets into two groups with diameter below (n=236) or above (n=195) 2 μm, we find that the chemical differences for the LD groups are statistically significant (*c*). Values are displayed as mean ± SD. Significance was determined using an unpaired t-test with Welch’s correction and p values are classed as *** <0.001, **** < 0.0001.

## Discussion

Lipid depots are present in many species, and their neutral lipid composition can be linked to certain diseases or is a potential engineering target for bioenergy production. Almost all studies focused on the changes in LD TAG composition have been based on data collected from large cell populations and thus only few insights into LD heterogeneity have been made (53). In this study, we present a method to extract quantitative information about spatial heterogeneity in LD chemistry, in terms of TAG chain length and double bonds with sub-micrometer resolution from non-invasive, broadband CARS imaging. While mass spectrometry can deliver complete chemical identification and micron-scale spatial resolution, it relies on extensive sample preparation, such as lipid extraction or embedding in complex matrix. Moreover, large and clean samples are required, which are not always available. Using a non-invasive optical method allows analysis of individual LDs in single cells as well as standard oils within minutes. Another feature of the BCARS method shown here is the ~ 2-fold better spatial resolution (46) compared to state-of-the-art imaging mass spectrometry (>1.4 μm resolution (31)). Principally, the resolution of our method is limited by diffraction to ~ 400 nm with better quality objectives. Finally, in contrast to previous studies employing spectral CARS (37, 38) our method does not return ratiometric quantities of TAGs but rather physiologically relevant numbers.

### Method reliability

Standard TAGs were used to establish the training matrix needed to generate the lipid components for the known covariance least square fitting, and food oils were subsequently used for the estimation of the standard error against industry-standard GC-MS. The error in our method is highest for lower chain lengths and higher degrees of saturation, *i.e.*, lower number of C=C. Physiologically relevant TAGs with chains length from 16 – 18 C and 1 – 2 C=C bonds can be determined with high accuracy. Specifically, for chains lengths below 16 the root mean square (RMS) deviation relative to the average chain length is approximately 6.4% while for chain lengths above 16 the RMS deviation is 3.7%. For oils with 0-1 double bonds on average, the relative deviation is 33.1 %, while the metric is 10.6 % for > 1 C=C bond. The error is evenly distributed around the GC-MS values, suggesting that there is no systematic error, *e.g.*, by an insufficient training matrix, causing these errors. Instead the deviation can be viewed as the true error of the method and in repeatability of the acquired data, which was taken over more than one year on a multi-user BCARS microscope.

### *In situ* lipid mapping

When applying the technique to more complex systems, like cells, other aspects must be considered for the spectral decomposition. If the cell had a lipid droplet size approaching or below our focal volume, as was the case for algae and yeast cells, it was necessary to include a protein and dilution component in the fitting since individual pixels in the lipid-rich regions will undoubtedly contain mixed cytosol and lipid signals in the spectral response. Yeast and algae with LDs < 0.5 μm are clear examples where the technique meets its limitations as no individual LDs can be seen with the BCARS spectral imaging. Nevertheless we can still use this method to compare lipid biochemistry within and among individual cells.

Although lipid droplets in microalgae are to date not well understood (54), it has been found that the cellular lipid storages in *Nannochloropsis* increase, in size and number, under nitrogen depletion (55). It has also been shown that the same strain of microalgae grown under nitrogen starvation contains around 15.0±1.3% saturated fatty acids and 21.2±1.7 % unsaturated fatty acids (45). This results in an average chain length of 16.9 with 1.4 double bonds. The batch we analyzed similarly shows high unsaturation (0.94 C=C).

For *yarrowia lipolytica*, where each cell has a > 500 nm TAG-filled LD and differentiated 3T3-L1, a model for mammalian adipocytes, we found good agreement among BCARS imaging, GC-MS measurements, and existing literature (56). However, the GC-MS method requires extensive sample preparation: TAG extraction from millions of cells (Supplemental Methods S1), while the BCARS method can be applied on as few as 5-10 cells. Including the time for image acquisition and image fitting (on a standard desktop PC), results can be within 10 hours. Interestingly, when looking within single LDs, the adipocytes show the saturated TAGs located on the outer parts of the droplet, almost the exact opposite what was seen for algae (and yeast). When comparing among LDs within adipocytes, we found that the unsaturation of TAGs inversely correlates with the size of the LDs, with large LDs having 11% lower # of C=C bonds than small LDs on average. During differentiation from mesenchymal precursors into adipocytes, LD sizes typically increase while LD number per cell decreases. It has previously been suggested that the size of the LDs in 3T3-L1s reflect the “age” of the droplet in that a small droplet is recently formed and that large droplets are older (57). Combined with the fact that lipid profiles of 3T3-L1s change during differentiation – with longer chains and more unsaturated fatty acids diminishing during the differentiation (58), this supports our finding that large (old) LDs have decreased chain length and unsaturation compared to small (young) LDs.

## Conclusion

In this work we have developed a method to generate fast overview of average lipid species, *in situ*, in mammalian and non-mammalian cells with up to 500 nm spatial resolution. This methodology allows for a somewhat detailed understanding of intra-LD, inter-LD, and cell-to-cell TAG chemical variation with virtually no sample preparation. Physiologically meaningful lipid quantification, in contrast to ratiometric quantities, is achieved, which shows high accordance with GC-MS-derived TAG chain length and saturation for pure oils. Due to the fast image acquisition and noninvasive nature of our method, future applications to study dynamic lipid biochemistry in cells, tissues, or in food emulsions will be enabled with much improved accuracy due to the straightforward *in situ* measurement conditions.

## Author contributions

A.P. and S.H.P conceived the idea, designed the project, and wrote the majority of the manuscript. A.P. collected the majority of data and analyzed all data, Y.W. contributed spectra of diluted oils, S.K. and C.B. contributed to the cell culture. C.B. and M.B. commented and edited the text. All authors read and approved the final version of the text.

## Acknowledgments/grant support

The authors thank Charlotta Olofsson, Gothenburg University, for the provision of 3T3-L1 cells; David Bergenholm and Jens Nielsen, Chalmers, for *Saccharomyces cerevisiae*; Oliver Konzock, Chalmers, for *yarrowia lipolytica*; Joshua J. Mayers, Chalmers, for *Nannochloropsis* sp; Mischa Schwendy, MPIP, for RAW and HEK cells; Frederik Fleissner and Marc-Jan van Zadel, MPIP, for support with the BCARS setup; Xiao Ling, MPIP, for input to the global fitting; and Pernilla Wittung-Stafshede, Chalmers, for great discussion. The research leading to these results has received funding from the European Union’s Seventh Framework Program (FP7/2007-2013) under grant agreement n°607842, Kungl. Vetenskaps-och Vitterhets-Samhället, Wilhem och Martina Lundgrens Stiftelse, and Marie Curie Foundation n°CIG322284.

